# Multi-genome synteny detection using minimizer graph mappings

**DOI:** 10.1101/2024.02.07.579356

**Authors:** Lauren Coombe, Parham Kazemi, Johnathan Wong, Inanc Birol, René L. Warren

## Abstract

In recent years, the landscape of reference-grade genome assemblies has seen substantial diversification. With such rich data, there is pressing demand for robust tools for scalable, multi-species comparative genomics analyses, including detecting genome synteny, which informs on the sequence conservation between genomes and contributes crucial insights into species evolution. Here, we introduce ntSynt, a scalable utility for computing large-scale multi-genome synteny blocks using a minimizer graph-based approach. Through extensive testing utilizing multiple ∼3 Gbp genomes, we demonstrate how ntSynt produces synteny blocks with coverages between 79–100% in at most 2h using 34 GB of memory, even for genomes with appreciable (>15%) sequence divergence. Compared to existing state-of-the-art methodologies, ntSynt offers enhanced flexibility to diverse input genome sequences and synteny block granularity. We expect the macrosyntenic genome analyses facilitated by ntSynt will have broad utility in generating critical evolutionary insights within and between species across the tree of life.

## Main

Rapid advances in both genome sequencing technologies and assembly algorithms have led to an explosion of genome sequences assembled at chromosome scale^1^. From pangenomics initiatives, such as the Human Pangenome Project^2^, to programs assembling a wide variety of non-model organisms, like the Earth BioGenome Project^3^, researchers now have unprecedented access to high-quality reference genomes. Robust bioinformatics tools are crucial to leverage this influx of data for comparative genomics studies, which can enable important insights into genomic influences on phenotypes, genomic diversity, and genome synteny^4–7^.

Genome synteny studies center on analyzing the conservation of genome structure within or between species^8–11^. The analysis can focus on large-scale macrosynteny, which tolerates smaller rearrangements within synteny blocks, or more fine-grained microsynteny^12–14^. Assessing synteny between genomes enables deeper insights into the evolution of genome structure, and supports a deeper functional understanding of genomes^15^. Synteny blocks can be broken at various types of genomic rearrangements, including inversions, translocations, insertions, and deletions^8,16^.

Studies of genomic synteny, such as research by Nadeau and Taylor^17^ into the conservation between human and mouse genomes in the 1980s, predates the sequence assembly of large genomes. Many early synteny studies relied on the labour-intensive and low-resolution laboratory technique of chromosome painting^18^, which involves hybridizing chromosome-specific fluorescent probes to cytogenetic slides. Starting in the 2000s, the increased availability of genomic resources lead to the development of multiple anchor-based bioinformatic synteny detection tools, including DAGchainer^15^, SynChro^9^, MCScanX^19^ and DRIMM-Synteny^20^. Each of these tools detect the conserved order of anchors or markers^8,21^, generally homologous genes, to detect synteny, although DRIMM-Synteny can use anchors based on other definitions of similarity. SynChro uses reciprocal best hits from the input genes to construct synteny backbones, and MCScanX chains collinear blocks based on BLASTP^22^ outputs using a dynamic programming approach. DAGChainer and DRIMM-Synteny both use graph-based approaches, with DAGChainer traversing a directed acyclic graph, and DRIMM-Synteny using an A-Bruijn graph-based representation.

More recently, improvements in sequence mapping and alignment algorithms have enabled tools to detect conserved stretches of DNA without requiring gene annotations. By using a whole genome approach, as opposed to sparse anchors, they compute synteny between the entire input genome sequences. Satsuma^23^, SyMap^14^, SyRI^7^ and halSynteny^13^ compute synteny blocks on pairs of input genomes. Satsuma aligns the input genomes using a Fast Fourier transform approach, while the others require sequence alignments as inputs. SyMap uses MUMmer^24^ alignments, which are clustered into anchors and chained using dynamic programming. SyRI is designed to compare chromosome-level genome assemblies, and utilizes minimap2^25^ or MUMmer alignments. While Progressive Cactus^26^, the aligner recommended for halSynteny, can compute multi-genome sequence alignments, halSynteny itself compares pairs of genomes. When computing multi-genome synteny blocks, SibeliaZ^27^ and SyntenyPortal^10^ are two state-of-the-art utilities. SibeliaZ computes multi-genome synteny blocks using compacted de Bruijn graphs, with graph traversal using carrying paths. With this approach, SibeliaZ is suited to computing locally collinear synteny blocks between multiple similar genomes, such as those belonging to different strains of a species. Finally, SyntenyPortal is a web application that computes synteny blocks between genome assembly builds from the UCSC Genome Browser^28^. SyntenyPortal utilizes pre-computed pairwise alignments from this database with inferCars^29^ to determine synteny block coordinates at four pre-specified resolutions. Therefore, none of these tools offer flexible and customizable computation of large-scale synteny blocks on multiple genomes.

Today, an increasing number of bioinformatics tools are utilizing minimizer sketches^30^, which represent underlying sequences using a particular subset of *k*-mers (substrings of length *k*) for various applications, including mapping^25^, estimating sequence divergence^31^, and assembly scaffolding^32,33^. Sketching greatly reduces the computational cost of comparing and analyzing sequences, making it an attractive approach for scalable tool development in support of large-scale genomics research.

In this work, we adapt the minimizer graph introduced in ntJoin^32^ for analyzing genome synteny. ntJoin^32^ utilizes undirected minimizer graph sketches to map genome sequences to one another for reference-guided scaffolding.

Here, we introduce ntSynt, a scalable utility for computing large-scale multi-genome synteny blocks. ntSynt uses lightweight, Bloom filter^34^-guided minimizer sketches to create an undirected minimizer graph, which is then leveraged for synteny block computation (Fig. 1, Supplementary Fig. 1). After graph simplification and breaking synteny blocks at putative large indels, the graph is extended with increasingly dense minimizer sketches to enhance the resolution of the synteny block coordinates. Finally, collinear blocks are merged to output the final synteny blocks. Each step of the ntSynt pipeline is flexible to user input genome sequences (assemblies) and parameters, making ntSynt a broadly useful utility. We show how ntSynt produces contiguous and accurate synteny blocks for genomes of increasing divergences, relatively quickly and with a low memory footprint, enabling a vast array of comparative genomics study designs.

**Fig. 1:**
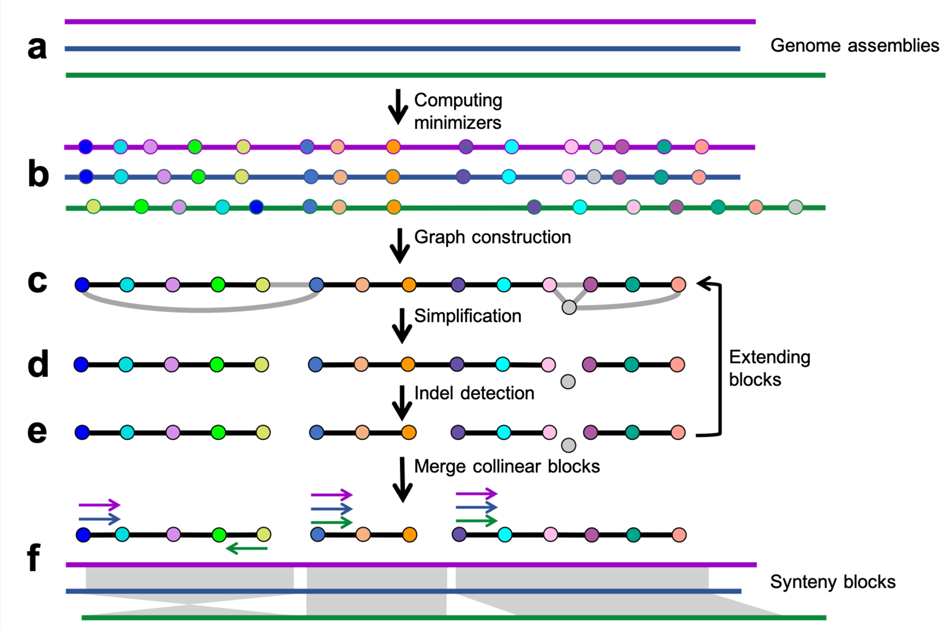
Schematic of the ntSynt workflow. (a) Multiple genome assemblies are provided as input to ntSynt, represented here with purple, blue and green lines. (b) First, minimizer sketches are computed from each input genome assembly using the *indexlr* functionality of btllib^43^ with a Bloom filter comprised of the *k-*mers common to all assemblies. The minimizers are shown as circles, with the same fill colour indicating identical minimizers. (c) Next, the minimizers that are found in all assemblies and are unique in each individual assembly are used to create an undirected minimizer graph, where the nodes are minimizers and edges are created between adjacent minimizers. The edge weights correspond to the number of genome assemblies that support the edge (represented as black for full assembly support and grey for partial assembly support, respectively). (d) The graph is simplified and edges are filtered to produce a series of linear graphs, or paths, that correspond to the initial synteny blocks. (e) The paths are broken at putative indels, and the graph can be extended to increase the block resolution using higher density minimizer sketches (arrow to (c)). (f) Finally, collinear blocks are merged, and the final synteny blocks are output. The coloured arrows indicate the relative orientation of the synteny blocks for each genome assembly. Synteny blocks between genomes are shown using the grey ribbons, with direct synteny represented by rectangles and inverted synteny between the blue and green sequences represented by the crossing ribbon.

## Results

### Pairwise synteny block detection using a simulated human genome

We first evaluated ntSynt and three comparators (SibeliaZ, halSynteny, SyRI) by generating pairwise synteny blocks between the human reference genome^35^ and a simulated rearranged human genome, with SNVs (single nucleotide variants) and indels (small insertions/deletions) introduced at various rates^36,37^ (Fig. 2a-d, Supplementary Tables 1-2). Compared to SibeliaZ, ntSynt generates substantially more contiguous synteny blocks, with block NG50 lengths (at least half of the genome covered by blocks ≥ this length) 1,525 to 9,514 times higher across the tested SNV and indel (variant) rates. In addition, the ntSynt synteny blocks have higher coverage, from 10.2–19.9% higher coverages for the lowest (0.1%) and highest (5.5%) variant rates, respectively.

**Fig. 2:**
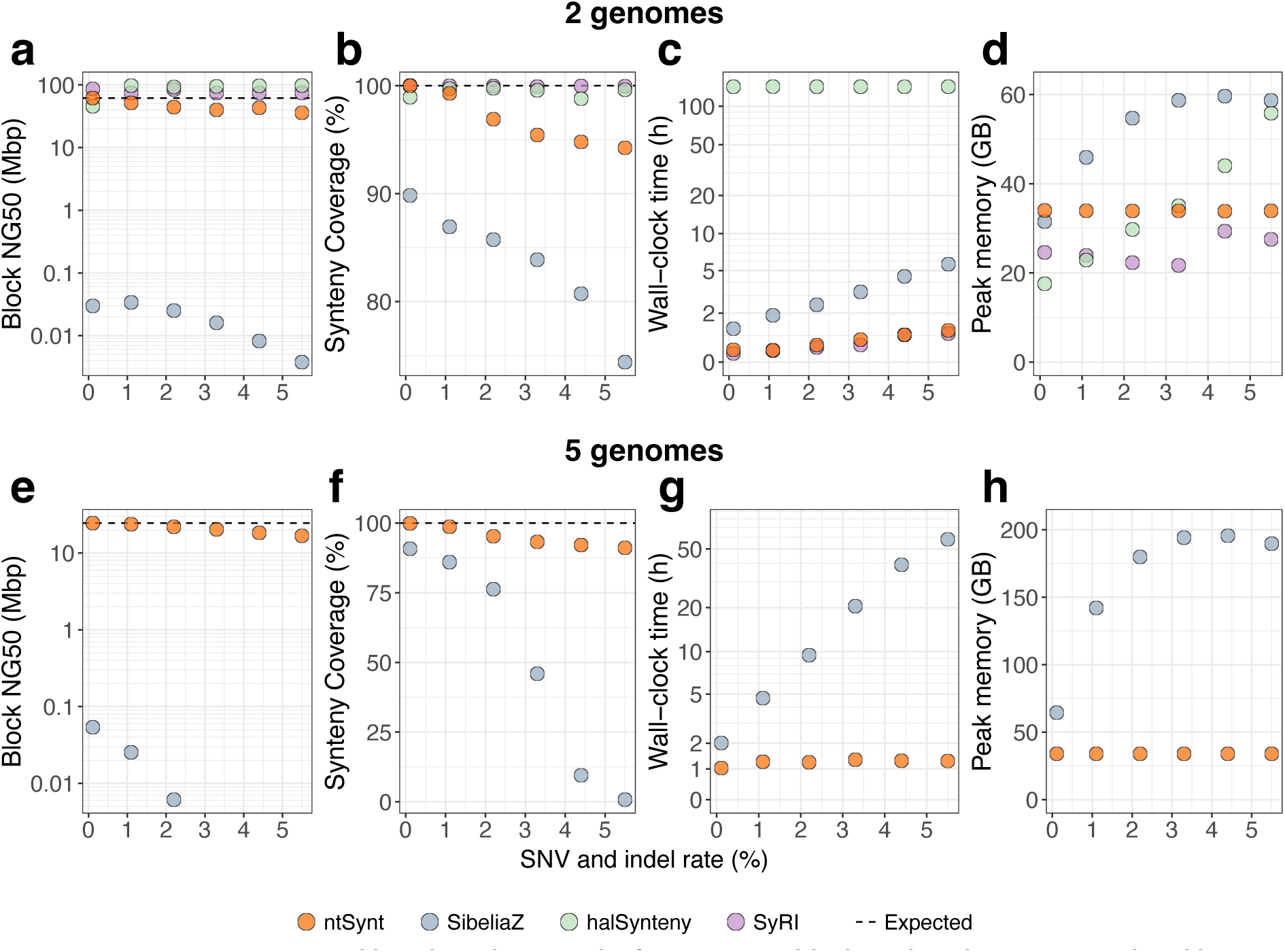
Contiguity, coverage and benchmarking results from synteny block analysis between simulated human genome sequences. Rearrangements in the human genome were simulated using SURVIVOR^36^, and SNVs and indels were introduced at increasing rates (ranging from 0.1% to 5.5%) using pIRS^37^. The top row plots (a-d) show the results of the pairwise comparisons between the human reference genome (T2T) and 1 rearranged human genome, while the bottom row (e-h) shows the results of a multi-genome synteny comparison between the human reference genome and 4 rearranged human genomes. halSynteny (green) and SyRI (purple) are only shown in a-d, as they are pairwise synteny utilities, while the multi-genome synteny tools ntSynt (orange) and SibeliaZ (blue) are shown in all plots. The synteny block NG50 length and synteny coverage statistics (a, b, e, f) were averaged over all input genomes. The horizontal dashed lines represent the expected value for the corresponding statistic based on the ground truth. Plots a,c,e,g are shown in log-linear scale, while b,d,f,h are in linear scale. Benchmarks were averaged over triplicate runs, and the average wall-clock time and peak memory values plotted.

Compared to the pairwise synteny detection tools tested, ntSynt synteny blocks generally have lower block NG50 lengths, though these contiguities do not exceed the expected NG50 length of 61.7 Mbp given the ground truth (35.9–61.7 Mbp). SyRI and halSynteny behave differently, with 74.3–86.3 Mbp and 92.6–97.7 Mbp block NG50 lengths, respectively. The synteny coverages for SyRI and halSynteny are over 98% for all tests, while the ntSynt coverages are over 99% for lower variant rates (0.1–1.1%), but 94–97% for the more divergent tests. Genomic regions that are not covered by ntSynt synteny blocks in these runs are mainly (>95%) centromeric (Supplementary Table 3).

In these pairwise tests, ntSynt ran 2.9 to 4.7-fold and 118.2 to 318.1-fold faster than SibeliaZ and halSynteny, respectively. SyRI was marginally (1.1–1.4 times) faster than ntSynt, both tools running in less than 1.5h across all variant rates tested. ntSynt and SyRI had fairly steady RAM usage (33.6–34.1 GB and 21.7–29.4 GB, respectively), but the RAM usage of SibeliaZ and halSynteny increased with higher variant rates, peaking at 59.6 GB and 60.9 GB, respectively.

### Synteny blocks between multiple simulated human genome sequences

Next, we computed synteny blocks between multiple simulated input genomes with ntSynt and SibeliaZ (Fig. 2e-h, Supplementary Tables 4-5). Five genomes were compared: the human reference genome and four genomes with simulated rearrangements. In all runs, ntSynt generated synteny coverages between 91.1–99.9%, and block NG50 lengths between 16.8–24.6 Mbp, approaching the ground truth block NG50 length (24.6 Mbp). Similar to the pairwise comparisons aforementioned, ntSynt generates synteny blocks with block NG50 lengths 457 to 3,584-fold longer than SibeliaZ, and its synteny coverage was 9.1–90.8% higher. For variant rates above 3%, the SibeliaZ synteny blocks have less than 50% genome coverage, thus the associated block NG50 lengths could not be computed. Furthermore, in these tests ntSynt runs faster and has a lower memory footprint than SibeliaZ, with runtimes and peak memory usages ranging from 50m to 2h and 33.9 to 34.1 GB of RAM for ntSynt, in contrast to 1.7h to 72.2h and 64.4 to 195.6 GB of RAM for SibeliaZ.

### Computing synteny blocks between four primate reference genomes

We computed synteny blocks between four primate reference genomes belonging to human, chimpanzee, bonobo, and gorilla with ntSynt and SibeliaZ (Fig. 3a, Supplementary Tables 6-7). Consistent with our simulated genome tests, ntSynt generated larger synteny blocks than SibeliaZ, with ntSynt blocks having an NG50 length of 7.6 Mbp, compared to 52.5 kbp for SibeliaZ. ntSynt also achieved higher synteny block coverage (92.2% vs. 88.6% for ntSynt and SibeliaZ, respectively).

**Fig. 3:**
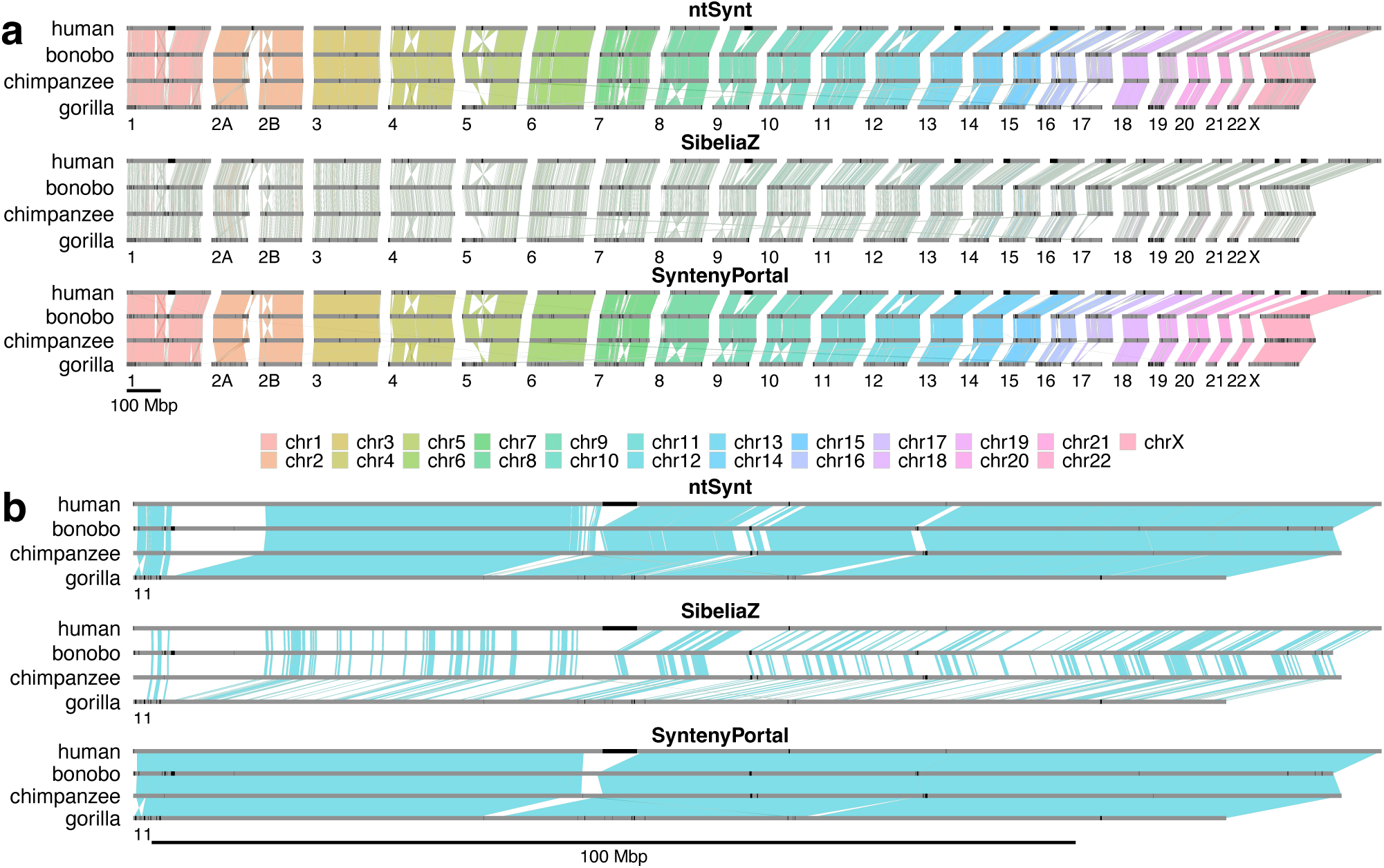
Ribbon plots showing synteny blocks computed between human, bonobo, chimpanzee and gorilla genome sequences using ntSynt, SibeliaZ and SyntenyPortal. The chromosomes are indicated using grey bars, with gaps and centromeres marked in black. Each ribbon is based on a synteny block >= 150 kbp that includes all 4 genomes. The ribbon colours represent the chromosome of the human genome in the synteny block, and each ribbon has a light grey border. The numbers below each plot indicate the chromosome in the other primate genomes, and the scales are indicated by the bar under each combined plot. Panel (a) shows the synteny blocks for the full genomes, and panel (b) zooms into chromosome 11. The ribbon plots were generated using the gggenomes R package (version 0.9.12.9000)^46^. Twisted ribbons depict inverted sequence synteny while direct synteny blocks are represented by parallelograms.

The outputs of ntSynt and SibeliaZ were also compared with synteny blocks from the SyntenyPortal web application (Fig. 3a). The synteny blocks computed using each approach are largely congruent, as evidenced by the consistent ribbon colours and transitions between the genome sequences for each chromosome. Although SyntenyPortal generates more contiguous synteny blocks than ntSynt (48.1 Mbp vs. 7.6 Mbp block NG50 lengths, respectively), this resource misses large variations between the genomes, as highlighted in Fig. 3b. Here, the ntSynt and SibeliaZ plots reveal a large (∼10 Mbp) deletion around the 4 Mbp position of chromosome 11 in the gorilla genome. However, a single SyntenyPortal synteny block spans this large deletion. This deletion was recapitulated using pairwise minimap2^25^ alignments (Supplementary Fig. 2a). Aligning gorilla long reads to the human reference genome (GRCh38) showed a median coverage of 14 over the region identified by the synteny block analysis as a putative deletion in gorilla (Supplementary Table 8). In this chromosome, there are additional examples of large indels missed by SyntenyPortal but detected using ntSynt (Supplementary Fig. 2a). The chromosome 11 SyntenyPortal blocks are only broken at a ∼2 Mbp insertion in the gorilla genome, a rearrangement that ntSynt also detected (Supplementary Fig. 2b).

### Computing synteny blocks between four primate genome assemblies

In a separate experiment, synteny blocks were computed between GoldRush^38^ and hifiasm^39^ human, bonobo, chimpanzee and gorilla genome sequence assembly drafts using ntSynt and SibeliaZ (Supplementary Tables 9-10), the only two utilities that can handle the simultaneous comparison of more than two genomes. The ntSynt four-genome synteny blocks had a coverage of 82.8% with 1.3 Mbp block NG50, while SibeliaZ generated synteny blocks with 79.1% coverage and 54.7 kbp block NG50 length.

### Assessing synteny between human, mouse and rat reference genomes

Finally, synteny blocks were computed between human, mouse, and rat reference genomes using ntSynt and SyntenyPortal (Fig. 4, Supplementary Tables 6 and 11). As SibeliaZ, which is only recommended for comparing closely-related genomes^27^, yielded synteny blocks with less than 3% coverage for this experiment, it was omitted from the analysis. These genomes have high sequence divergence, with Mash^31^ estimating pairwise sequence divergences up to 18.8%. The ntSynt synteny blocks between the three genomes had 79.2% coverage and 3.9 Mbp block NG50 length, while SyntenyPortal generated blocks with 89.8% coverage and 7.1 Mbp block NG50. As evident from the chromosome sequence painting plots (Fig. 4), the synteny blocks computed by ntSynt and SyntenyPortal are largely consistent in terms of chromosome identities and block orientations. Notably, the majority of the large gaps seen in the ntSynt synteny blocks intersect with centromeric and gap sequences, as evidenced in human chromosomes 1, 4, 5, 11 and 12. ntSynt generated synteny blocks between these three divergent genomes in 1.1h using 34.2 GB of RAM (Supplementary Tables 11-12).

**Fig. 4:**
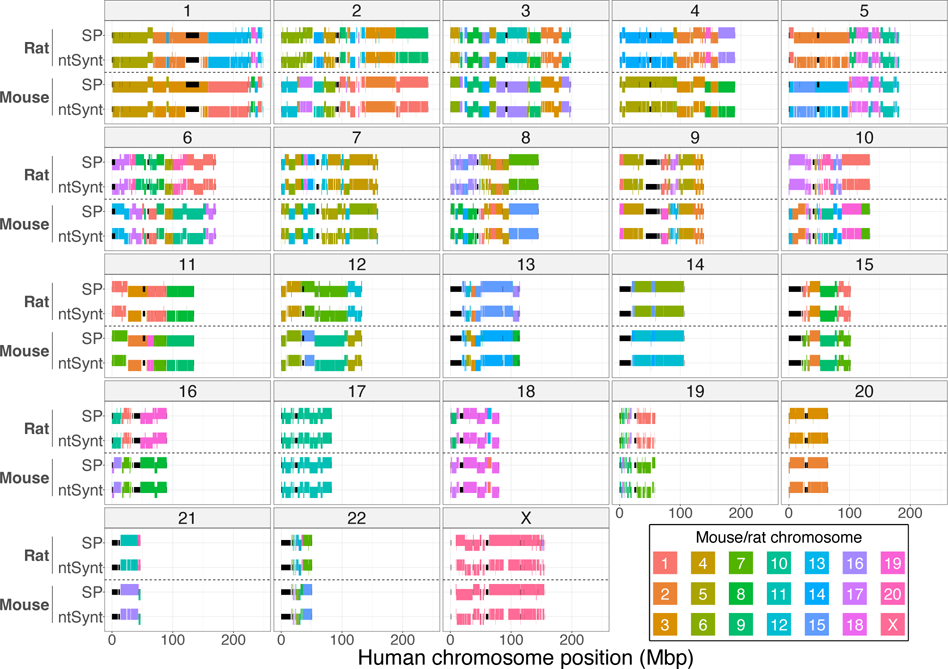
Chromosome sequence painting plots comparing synteny blocks between human, mouse, and rat reference genomes computed by ntSynt and SyntenyPortal (SP). Each facet is labeled with the corresponding human chromosome. The segments are coloured by the chromosome in mouse or rat, respectively. Each segment represents a 3-genome synteny block in ntSynt or SyntenyPortal. The segments being nudged vertically up or down represent the forward or reverse orientation, respectively. The smaller black segments indicate gaps and centromere sequence in the human reference genome build.

## Discussion

ntSynt introduces a novel approach for detecting multi-genome synteny blocks by employing minimizer graphs to simultaneously map multiple genome sequences to one another. Unlike anchor-based synteny detection tools, ntSynt does not require annotations or precomputed markers, and utilizes full input genome sequences. With the introduction of various graph simplification and traversal algorithms, ntSynt builds upon the undirected minimizer graph approach introduced by ntJoin^32^, adapting the principles for genome synteny detection. ntSynt also introduces a minimizer selection approach guided by a common Bloom filter^34^, a probabilistic data structure used to store the *k*-mers found in all input genome assemblies. These algorithms enable ntSynt’s robustness to complex genome graphs without sacrificing the sensitivity of the multi-genome mappings. The effectiveness of this approach is demonstrated by the resulting megabase-range synteny block NG50 lengths with coverages over 92% when comparing four primate reference genomes. Moreover, ntSynt’s promise in supporting genome evolution studies is highlighted by the identification of multiple known rearrangements between primate genomes, such as the fusion of chromosome 2 in human^40^, and a ∼47 Mbp inversion in human chromosome 12^41^ (Fig. 3).

Utilizing undirected minimizer graphs for multi-genome mapping, combined with the graph traversal and post-processing algorithms, allows for minor rearrangements and base-level differences between the input genome sequences to be tolerated within a synteny block, and substantial flexibility in the granularity of the ntSynt macrosynteny blocks. The construction and traversal of these graphs are controlled by multiple parameters, including the merge, indel, and block size thresholds. These parameters have empirically determined default values based on a specified genome sequence divergence, but can be fine-tuned to refine the synteny blocks for particular research questions. Generally, setting these thresholds higher allows for more contiguous synteny blocks and a broader view of the genome synteny. Conversely, lowering the thresholds will give a more granular picture of the genome synteny, making these parameters a balance between synteny block resolution and output block contiguity. These variable criteria for defining synteny blocks highlight why the block NG50 length alone is not a perfect assessment statistic.

The other synteny block detection utilities tested have more limited parameters, constraining their flexibility in achieving these different resolutions. The minimum block size can be specified for SibeliaZ, halSynteny and SyntenyPortal, but only the pairwise comparators halSynteny and SyRI have parameters for controlling the tolerated distance between anchoring alignments in a synteny block. None of the comparator tools have clear parameters controlling indel detection.

In addition to tunable parameters, ntSynt is also flexible to the number, divergence and contiguity of the input genomes. ntSynt computes synteny blocks with megabase-scale contiguity and high coverage for closely-related genomes, such as those of primates, and more divergent genomes, like the human-mouse-rat tests. This robustness to increasing divergence was also demonstrated by achieving synteny coverages above 91% when comparing five simulated human genomes, each with up to 5.5% variant rates between them. This also demonstrates that using a mapping-based approach for synteny analysis, versus an annotation-guided anchor-based approach, is effective for more divergent genomes, despite the contrary speculation reported in previous studies^8^. The robustness of the paradigm is further supported by the high multi-genome synteny block coverage (>82%) reached by ntSynt when comparing genome assemblies of human (assembled with GoldRush^38^), bonobo, chimpanzee and gorilla (assembled with hifiasm^39^).

While halSynteny and SyRI produced synteny blocks with high coverage (>98%) in the simulated genome tests, they are limited to pairwise genome comparisons. By design, SyRI utilizes the pairwise sequence aligner minimap2^25^ to generate alignments for computing synteny blocks. Progressive Cactus^26^, the alignment engine behind halSynteny, did not scale well for the pairwise human comparisons, each test requiring around 6 days to complete. Finally, both tools generate synteny blocks with higher than expected NG50 lengths, indicating that they do not break the blocks at all ground truth rearrangements.

Although SibeliaZ and SyntenyPortal compute multi-genome synteny blocks, they are less adaptable than ntSynt in the variety of compatible input genomes. SibeliaZ is designed for identifying locally collinear genome segments in closely-related genomes, or those with an evolutionary distance of less than 0.09 substitutions per site to the most recent common ancestor^27^. This restriction leads to lower synteny coverage as the genomic sequence divergence increases, especially when comparing multiple genomes, as demonstrated in the simulated tests, and meant that SibeliaZ was not suitable for our human-mouse-rat experiment due to the high sequence divergence between those species’ genomes (up to 18%). SibeliaZ’s microsynteny focus was also evident from the contiguity of the synteny blocks, as ntSynt generated 1,400-times higher block NG50 lengths on average. On the other hand, SyntenyPortal can compare more divergent genomes, as seen by the >89% coverage when comparing human, mouse and rat genomes, but can only use genome builds that are available on the UCSC genome browser^28^. Thus, we could not use SyntenyPortal in our simulated tests, nor for comparing the GoldRush and hifiasm genome assemblies. While both SyntenyPortal and SibeliaZ are restricted in the synteny analyses that they can be used for, ntSynt can compare genome assemblies with a range of divergences.

In addition to its inflexibility towards the input genome assemblies, SyntenyPortal fails to break synteny blocks at various large indels. Examples can be seen in Fig. 3b, where multiple large indels were missed by SyntenyPortal, but detected by ntSynt, including a ∼10 Mbp deletion in chromosome 11 of the gorilla genome. Further investigation into this large deletion using gorilla long sequencing reads suggested that the deletion is likely due to a misassembly in the older gorilla assembly release used herein. ntSynt breaking the synteny block at this region, and thus flagging it for further analysis, enabled the observation of this genome assembly issue. This feature may prove useful for evaluating the completeness and consistency of future genome assembly projects.

As well as its adaptability to input genome sequences and synteny block granularity, ntSynt is also relatively fast and memory-efficient, largely due to its graph-based design and use of sequence minimizers. In all tests, comparing up to five ∼3 Gbp genomes, ntSynt required at most 2h and 34.2 GB of RAM. In comparison, SibeliaZ, the only command-line multi-genome synteny comparator, was more computationally expensive as the number and divergence of compared genomes increased, requiring at most 72.2h and 195.6 GB RAM. Furthermore, the RAM usage of ntSynt remained between 33.6–34.2 GB in all tests, demonstrating that the required memory does not correlate with the number of input genomes. The most memory-intensive step was the creation of the common *k-*mer Bloom filter^34^, the probabilistic data structure we used to store the *k*-mers occurring in all input genome assemblies (Supplementary Table 12). The common Bloom filter employed by ntSynt is built using a cascading approach, which ensures that the peak memory usage is capped at twice the Bloom filter size. As this size is calculated based on the genome sizes and specified false positive rate, the memory can be further lowered by increasing the false positive rate. However, this is a balance, as increasing the false positive rate too aggressively can negatively impact the multi-genome mapping sensitivity.

Despite the contiguous and high coverage multi-genome synteny blocks achieved using ntSynt, the current implementation has a few limitations. Because only sequence minimizers that are unique in each individual assembly and found in all input genomes are integrated into the graph-based mapping, ntSynt will not detect duplications, which other tools, including SibeliaZ, can identify. Further, when comparing many highly divergent genomes, the requirement that only minimizers found in all input genomes are retained can result in a sparse graph and thus a lower synteny coverage. We do see a slight decrease in synteny coverage in our most divergent tests (e.g., human-mouse-rat) compared to the primate tests, although we also observe this decrease in the SyntenyPortal results. The impact of this issue could be mitigated by decreasing the *k*-mer size, which produces a denser minimizer sketch and minimizer graph (Supplementary Fig. 3).

As we have demonstrated, ntSynt enables the alignment-free, scalable and simultaneous computation of multi-genome synteny blocks for input genomes of various divergences. From deciphering pangenome structure within species to providing evolutionary insights between species, we expect the synteny blocks generated by ntSynt to lay the groundwork for many comparative genomics analyses, and enable the genomics community to more effectively leverage the continuously expanding diversity of genome assembly resources.

## Methods

ntSynt takes two or more genomes (reference or draft assembly grade sequences) in FASTA format as input, maps the input sequences to one another using minimizers, and outputs the synteny blocks in a BED-like format (Supplementary Fig. 1, Supplementary Table 13).

### Common Bloom filter construction

In the first step of the pipeline, ntSynt generates a common Bloom filter, which contains the *k*-mers found in all input genomes to be compared, using a multi-level cascading Bloom filter approach (Supplementary Figs. 1 and 4). Bloom filters are probabilistic data structures which represent the (probabilistic) presence or (deterministic) absence of elements using a bit array^34^. For *n* input assemblies, there will be *n* Bloom filters (BF) in the cascade, with at most two initialized and stored in memory at a given time. The size of each BF (*bfsize*, in bits) is determined using the input genome sizes and the specified false positive rate (*--fpr*, default 0.025) as shown in Equation 1, where *genome_size* is the largest input genome size in base pairs, and *fpr* is the false positive rate.

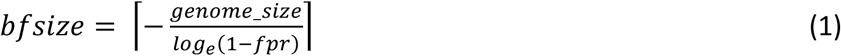

To initialize the cascade, the sequences in assembly *1* are *k*-merized using ntHash2^42^ and inserted into BF level *1*. Then, for each subsequent assembly *i* (2 ≤ *i* ≤ *n*), a new BF level *i* is initialized. If *i* > 2, the level *i* − 2 BF is deallocated prior to the level *i* BF initialization. The sequences in genome *i* are *k*-merized, and queried against BF level *i* − 1. The *k*-mers that are present in BF level *i* – 1 are inserted into BF level *i*. Once all input sequences have been *k*-merized and inserted into the corresponding BF level, the final BF level *n*, termed the *common Bloom filter*, is then used for computing minimizer sketches in subsequent steps.

### Constructing the initial undirected minimizer graph

Minimizer sketches are generated for each input sequence using the *indexlr* utility in btllib^43^, which generates minimizers based on the approach described in Roberts *et al*. 2004^30^. Briefly, for a given sequence, the canonical (reverse-complement invariant) hash values for *w* (window size, *-w*, default 1000) adjacent *k-*mers (substrings of length *k, -k*, default 24) are computed using ntHash2^42^. Only *k*-mers that are present in the common Bloom filter are considered when selecting the window’s minimizer. The *k*-mer with the smallest hash value is chosen as the minimizer for the given window, and the output minimizer hash value is computed using a second hash function. Minimizer sketches for all input genome sequences are generated by sliding the window across all sequences, and repeating the operation for each window, only adding to the sketch when there is a new minimizer. In addition to the minimizer hash value, *indexlr* also reports the position, strand and corresponding *k*-mer sequence for each minimizer.

The initial undirected minimizer graph is constructed as described in ntJoin^32^, where the nodes are minimizers, and edges are created between minimizers that are adjacent in at least one genome sequence input. The edge weights correspond to the number of input genomes in which the minimizers are adjacent (1 ≤ *edge-weight* ≤ *n*, where *n* is the number of input genomes). Only minimizers that are found in all input genomes and are unique in each individual genome will be retained and added to the minimizer graph.

### Filtering the minimizer graph

The minimizer graph is simplified by removing noisy minimizers that disrupt linear graph paths (Supplementary Fig. 5). First, we define fully anchored, partially anchored, and unanchored nodes. Fully anchored nodes have a degree of two and the sum of their incident edge weights is the maximum value (2*n*, where *n* is the number of input genomes). Partially anchored nodes have a degree of three, and one incident edge weight is *n*. All other nodes are considered unanchored. For each edge (*u, v*), where both *u* and *v* are partially anchored nodes, if there is an alternate path of length two (*u, z, v*) between the nodes, the middle node *z* is marked as noisy and removed from the graph. The weight of edge (*u, v*) is set as *n* after this removal. Following graph simplification, all remaining edges with a weight less than *n* are removed from the graph.

### Finding initial synteny blocks

The initial synteny blocks are identified from linear paths through the filtered minimizer graph. A given linear graph path can be converted to synteny block coordinates using the positions of the minimizers for each input sequence in that graph (Supplementary Fig. 6). Each linear path is first broken at edges which connect minimizers from different sub-sequences (or contigs) in any of the input genome sequences. Then, the start and end coordinates of the synteny block in each input sequence are computed from the terminal minimizers in the path. The orientation of the block in a given sequence input is determined by assessing if the minimizer positions in the graph are largely (≥ 90%) increasing or decreasing, which then assigns the orientation as “+” (forward) or “-” (reverse), respectively. If a block cannot be oriented, it is not included in the ntSynt output.

### Breaking synteny blocks at indels

To break the synteny blocks at indels, the interarrival distances between minimizers are assessed for each edge in the minimizer graph paths (Supplementary Fig. 7, Equation 2). For *n* input genomes, for each edge (*u, v*), where *a*_:.1J_ is the position of minimizer *a* in genome *x*, the indel score is computed as:

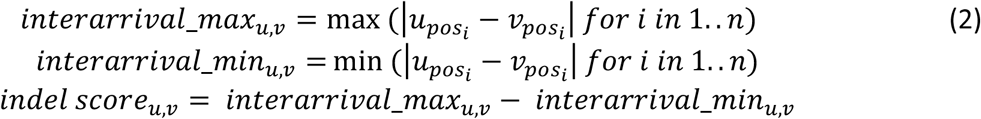

If the indel score for a given edge is greater than the specified indel score threshold (*--indel*), the edge is removed from the path, breaking the synteny block at the large indel.

### Dynamic minimizer graph extension with decreasing window size

The existing minimizer graph, or paths, can be extended using minimizers computed with decreasing window sizes (Supplementary Fig. 8). For a given window size *w_round* less than the original *w*, minimizers are computed (using the same *k-*mer size and common Bloom filter) on regions of each input genome sequence that are not covered by a synteny block. These sketches are used to dynamically extend the graph. Then, ntSynt repeats the graph simplification, filtering and indel detection steps to update the synteny blocks. This process of graph extension to increase block resolution using lower window sizes can be performed for any number of rounds (specified using a list of decreasing window sizes, *--w_rounds*).

### Merging collinear synteny blocks

Next, the resulting synteny blocks are sorted, filtered based on size (blocks greater than *-- block_size* are retained) and collinear synteny blocks separated by up to a threshold number of bases (*--merge*) are merged. Synteny blocks are considered collinear if they have consistent contig IDs, strands and positions (Supplementary Fig. 9), and are not separated by an indel.

### Final output synteny blocks

The final synteny blocks are output in a BED-like format, where each synteny block is assigned a unique block ID (Supplementary Table 13). Between each pair of synteny blocks, the final column specifies the reason for the discontinuity, which includes inconsistent contig IDs, orientations or positions, or exceeding the indel or merge thresholds.

### Implementation

The ntSynt pipeline is driven by a Snakemake^44^ file, which is launched using a python wrapper script. The common Bloom filter construction is implemented in C++, and all other algorithms are implemented in python. ntSynt is available from GitHub (https://github.com/bcgsc/ntSynt), and can be installed using conda. ntSynt requires the user to supply an estimated maximum sequence divergence (*--divergence %*) between the input genomes, which in turn selects presets with default settings for the *--block_size, --indel, --merge* and *--w_rounds* parameters (Supplementary Table 14). However, all parameters can be configured and independently fine-tuned by the user, and, when specified, override the presets selected by ntSynt in response to the input % divergence estimate.

### Evaluation

Four separate rearrangements of the human reference genome (T2T build) were simulated using SURVIVOR^36^ v1.0.7, with SNVs and indels introduced at various rates using pIRS^37^ v2.0.2. SURVIVOR was configured to simulate two translocations between 10–50 kbp, 20 inversions between 10–50 kbp and 20 indels between 50–100 kbp. For each rearranged genome sequence, SNVs were simulated at 0.1%, 1%–5% (step size 1), with indels simulated at 0.01%, 0.1–0.5% (step size 0.1), respectively. For pairwise synteny block detection, one rearranged human genome was compared to the human reference genome using ntSynt (v1.0.0), SibeliaZ^27^ (v1.2.5), halSynteny^13^ (v2.6.8) and SyRI^7^ (v1.6.3). As suggested in their documentation, SibeliaZ ran using the *-n* option to skip the alignment step, then the output was passed to maf2synteny^45^ (v1.2) to generate the final synteny blocks. The input pairwise alignments for halSynteny and SyRI were generated using Progressive Cactus^26^ (v2.6.8) and minimap2^25^ (v2.17-r941), respectively. The parameters used for each of these runs can be found in Supplementary Table 15. For multiple genome synteny block detection, the four rearranged human genomes with various SNV and indel rates and the human reference genome were compared using ntSynt and SibeliaZ (Supplementary Table 15). All runs involving simulated genome sequences were run in triplicate for benchmarking assessments.

To detect synteny blocks using multiple real genomes, we compared the human (GRCh38), bonobo (Mhudiblu_PPA_v0), chimpanzee (Clint_PTRv2) and gorilla (Kamilah_GGO_v0) reference genomes using ntSynt and SibeliaZ. Mash^31^ (v2.3, *-p* 12 *-s* 10000) was used to estimate the maximum sequence divergence between these input genomes, and the *-- divergence* parameter for ntSynt was subsequently set at 1.7. All other ntSynt parameters were kept at default settings, and SibeliaZ was run using the parameters listed in Supplementary Table 15. Synteny blocks between the same reference genome builds were also obtained using the SyntenyPortal^10^ web application (resolution=150 kbp). We compared the four-genome synteny blocks that were larger than 150 kbp using the gggenomes^46^ R package (v0.9.12.9000). To assess a large observed deletion in chromosome 11 of the gorilla genome, we aligned Pacific BioSciences (PacBio) HiFi reads (∼17-fold coverage) from the same gorilla individual (Kamilah) to the human genome reference (GRCh38) using minimap2 (Supplementary Table 8). We then assessed the read coverage over the region putatively deleted in gorilla using samtools^47^ (v1.16.1, samtools view chr11:4249395-14277464) and bedtools^48^ genomecov (v2.30.0). To further assess how these extracted reads aligned to the gorilla genome used herein as well as a newer gorilla genome assembly, we aligned the extracted reads (from samtools view) to each gorilla genome with minimap2, filtering for primary alignments with mapping quality ≥ 50 using samtools. The statistics of these alignments were assessed using bamtools^49^ (v2.5.1).

Using the same parameters as the previous primate reference genome runs, we also computed synteny blocks between *de novo* genome assemblies of the same primate species (human, bonobo, chimpanzee, gorilla) using ntSynt and SibeliaZ. The human genome assembly was generated using GoldRush^38^ with Oxford Nanopore long reads as input, while the other primate genome assemblies were generated using hifiasm^39^ with PacBio HiFi long reads as input (Supplementary Table 9).

Finally, synteny blocks were computed between human (GRCh38), mouse (GRCm39) and rat (Rnor_6.0) reference genome builds with ntSynt (Supplementary Table 6). The approximate sequence divergence rates between these genome assemblies were assessed using Mash, as described for the primate tests. Consequently, ntSynt was run with *--divergence* 18.8 *--indel* 500000, with all other parameters kept at the default values. Synteny blocks between these three genomes were also obtained from the SyntenyPortal web application (resolution=150 kbp).

All tests were run on a server with 144 Intel(R) Xeon(R) Gold 6254 CPU @ 3.1 GHz with 2.9 TB RAM using 12 threads. The python scripts used to assess the summary statistics of the output synteny blocks are available at https://github.com/bcgsc/ntSynt/tree/main/analysis_scripts, and example R scripts for generating the gggenomes^46^ ribbon plots and chromosome sequence painting plots are available at https://github.com/bcgsc/ntSynt/tree/main/visualization_scripts.

## Data availability

The accessions for all reference genome assemblies used in the described experiments can be found in Supplementary Tables 6 and 9. The simulated rearranged human genomes are available on Zenodo (https://doi.org/10.5281/zenodo.10627623).

## Code availability

ntSynt is freely available on GitHub (https://github.com/bcgsc/ntsynt).

## Acknowledgements

This study is supported by the Canadian Institutes of Health Research (CIHR) [PJT-183608, I.B.].

## References

1. Formenti, G. et al. The era of reference genomes in conservation genomics. Trends in Ecology & Evolution 37, 197–202 (2022).

2. Wang, T. et al. The Human Pangenome Project: a global resource to map genomic diversity. Nature 604, 437–446 (2022).

3. Lewin, H. A. et al. The earth BioGenome project 2020: Starting the clock. Proceedings of the National Academy of Sciences 119, e2115635118 (2022).

4. Lian, Q. et al. A pan-genome of 72 Arabidopsis thaliana accessions reveals a conserved genome structure throughout the global species range. PREPRINT (Version 1) available at Research Square (2023) doi:10.21203/rs.3.rs-2976609/v1.

5. Harling-Lee, J. D. et al. A graph-based approach for the visualisation and analysis of bacterial pangenomes. BMC Bioinformatics 23, 416 (2022).

6. Beier, S. & Thomson, N. R. Panakeia - a universal tool for bacterial pangenome analysis. BMC Genomics 23, 265 (2022).

7. Goel, M., Sun, H., Jiao, W.-B. & Schneeberger, K. SyRI: finding genomic rearrangements and local sequence differences from whole-genome assemblies. Genome Biology 20, 277 (2019).

8. Lallemand, T., Leduc, M., Landès, C., Rizzon, C. & Lerat, E. An Overview of Duplicated Gene Detection Methods: Why the Duplication Mechanism Has to Be Accounted for in Their Choice. Genes 11, (2020).

9. Drillon, G., Carbone, A. & Fischer, G. SynChro: A Fast and Easy Tool to Reconstruct and Visualize Synteny Blocks along Eukaryotic Chromosomes. PLOS ONE 9, e92621 (2014).

10. Lee, J. et al. Synteny Portal: a web-based application portal for synteny block analysis. Nucleic Acids Research 44, W35–W40 (2016).

11. Haug-Baltzell, A., Stephens, S. A., Davey, S., Scheidegger, C. E. & Lyons, E. SynMap2 and SynMap3D: web-based whole-genome synteny browsers. Bioinformatics 33, 2197–2198 (2017).

12. Robert, N. S. M., Sarigol, F., Zieger, E. & Simakov, O. SYNPHONI: scale-free and phylogeny-aware reconstruction of synteny conservation and transformation across animal genomes. Bioinformatics 38, 5434–5436 (2022).

13. Krasheninnikova, K. et al. halSynteny: a fast, easy-to-use conserved synteny block construction method for multiple whole-genome alignments. GigaScience 9, giaa047 (2020).

14. Soderlund, C., Bomhoff, M. & Nelson, W. M. SyMAP v3.4: a turnkey synteny system with application to plant genomes. Nucleic Acids Research 39, e68–e68 (2011).

15. Haas, B. J., Delcher, A. L., Wortman, J. R. & Salzberg, S. L. DAGchainer: a tool for mining segmental genome duplications and synteny. Bioinformatics 20, 3643–3646 (2004).

16. Vergara, I. A. & Chen, N. Large synteny blocks revealed between Caenorhabditis elegans and Caenorhabditis briggsae genomes using OrthoCluster. BMC Genomics 11, 516 (2010).

17. Nadeau, J. H. & Taylor, B. A. Lengths of chromosomal segments conserved since divergence of man and mouse. Proceedings of the National Academy of Sciences 81, 814–818 (1984).

18. Ried, T., Schröck, E., Ning, Y. & Wienberg, J. Chromosome painting: a useful art. Human Molecular Genetics 7, 1619–1626 (1998).

19. Wang, Y. et al. MCScanX: a toolkit for detection and evolutionary analysis of gene synteny and collinearity. Nucleic Acids Research 40, e49–e49 (2012).

20. Pham, S. K. & Pevzner, P. A. DRIMM-Synteny: decomposing genomes into evolutionary conserved segments. Bioinformatics 26, 2509–2516 (2010).

21. Liu, D., Hunt, M. & Tsai, I. J. Inferring synteny between genome assemblies: a systematic evaluation. BMC Bioinformatics 19, 26 (2018).

22. Altschul, S. F., Gish, W., Miller, W., Myers, E. W. & Lipman, D. J. Basic local alignment search tool. Journal of Molecular Biology 215, 403–410 (1990).

23. Grabherr, M. G. et al. Genome-wide synteny through highly sensitive sequence alignment: Satsuma. Bioinformatics 26, 1145–1151 (2010).

24. Marçais, G. et al. MUMmer4: A fast and versatile genome alignment system. PLOS Computational Biology 14, e1005944 (2018).

25. Li, H. Minimap2: pairwise alignment for nucleotide sequences. Bioinformatics 34, 3094–3100 (2018).

26. Armstrong, J. et al. Progressive Cactus is a multiple-genome aligner for the thousand-genome era. Nature 587, 246–251 (2020).

27. Minkin, I. & Medvedev, P. Scalable multiple whole-genome alignment and locally collinear block construction with SibeliaZ. Nature Communications 11, 6327 (2020).

28. Kent, W. J. et al. The human genome browser at UCSC. Genome Research 12, 996–1006 (2002).

29. Ma, J. et al. Reconstructing contiguous regions of an ancestral genome. Genome Research 16, 1557–1565 (2006).

30. Roberts, M., Hayes, W., Hunt, B. R., Mount, S. M. & Yorke, J. A. Reducing storage requirements for biological sequence comparison. Bioinformatics 20, 3363–3369 (2004).

31. Ondov, B. D. et al. Mash: fast genome and metagenome distance estimation using MinHash. Genome biology 17, 1–14 (2016).

32. Coombe, L., Nikolić, V., Chu, J., Birol, I. & Warren, R. L. ntJoin: Fast and lightweight assembly-guided scaffolding using minimizer graphs. Bioinformatics 36, 3885–3887 (2020).

33. Coombe, L., Warren, R. L., Wong, J., Nikolic, V. & Birol, I. ntLink: A Toolkit for De Novo Genome Assembly Scaffolding and Mapping Using Long Reads. Current Protocols 3, e733 (2023).

34. Bloom, B. H. Space/time trade-offs in hash coding with allowable errors. Communications of the ACM 13, 422–426 (1970).

35. Nurk, S. et al. The complete sequence of a human genome. Science 376, 44–53 (2022).

36. Jeffares, D. C. et al. Transient structural variations have strong effects on quantitative traits and reproductive isolation in fission yeast. Nature Communications 8, 14061 (2017).

37. Hu, X. et al. pIRS: Profile-based Illumina pair-end reads simulator. Bioinformatics 28, 1533–1535 (2012).

38. Wong, J. et al. Linear time complexity de novo long read genome assembly with GoldRush. Nature Communications 14, 2906 (2023).

39. Cheng, H., Concepcion, G. T., Feng, X., Zhang, H. & Li, H. Haplotype-resolved de novo assembly using phased assembly graphs with hifiasm. Nature Methods 18, 170–175 (2021).

40. Shao, Y. et al. Phylogenomic analyses provide insights into primate evolution. Science 380, 913–924 (2023).

41. Mao, Y. et al. A high-quality bonobo genome refines the analysis of hominid evolution. Nature 594, 77–81 (2021).

42. Kazemi, P. et al. ntHash2: recursive spaced seed hashing for nucleotide sequences. Bioinformatics 38, 4812–4813 (2022).

43. Nikolić, V. et al. btllib: A C++ library with Python interface for efficient genomic sequence processing. Journal of Open Source Software 7, 4720 (2022).

44. Mölder, F. et al. Sustainable data analysis with Snakemake [version 2; peer review: 2 approved]. F1000Research 10, (2021).

45. Kolmogorov, M. et al. Chromosome assembly of large and complex genomes using multiple references. Genome research 28, 1720–1732 (2018).

46. Hackl, T., Ankenbrand, M. J. & Adrichem, B. van. gggenomes: A Grammar of Graphics for Comparative Genomics. https://github.com/thackl/gggenomes (2023).

47. Danecek, P. et al. Twelve years of SAMtools and BCFtools. Gigascience 10, (2021).

48. Quinlan, A. R. & Hall, I. M. BEDTools: a flexible suite of utilities for comparing genomic features. Bioinformatics 26, 841–842 (2010).

49. Barnett, D. W., Garrison, E. K., Quinlan, A. R., Strömberg, M. P. & Marth, G. T. BamTools: a C++ API and toolkit for analyzing and managing BAM files. Bioinformatics 27, 1691–1692 (2011).

