## Supplementary Material for "Multi-genome synteny detection using minimizer graph mappings"

### Supplementary Information for: “Multi-genome synteny detection using minimizer graph mappings”

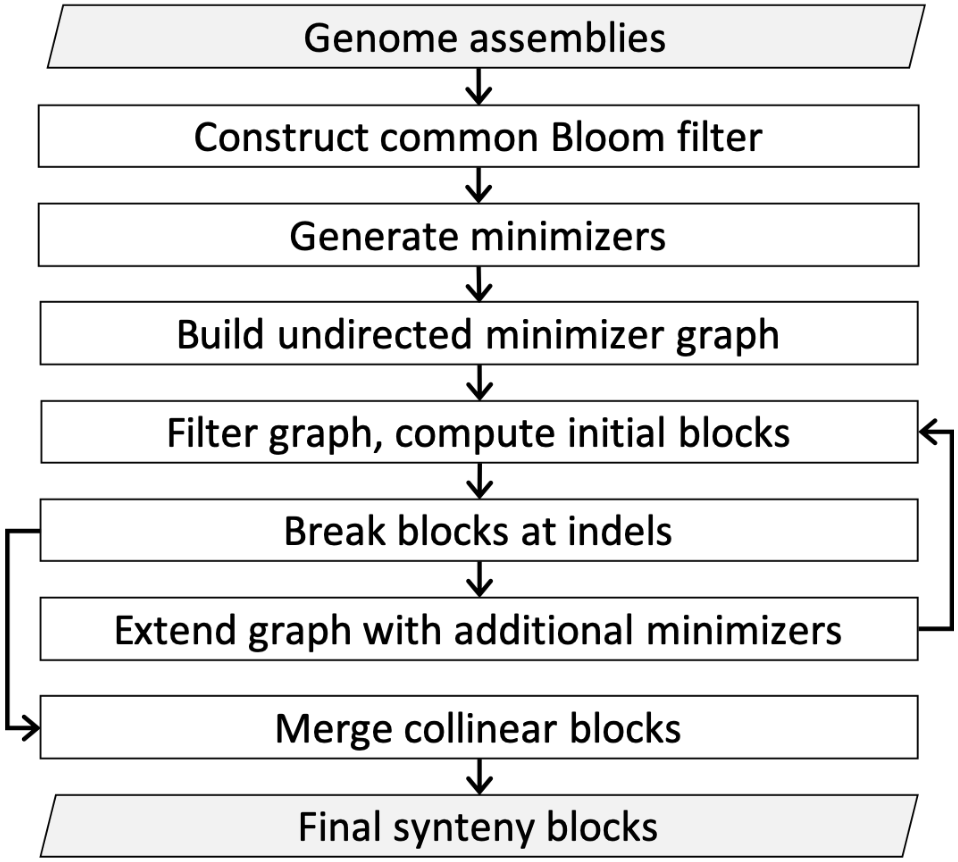

**Supplementary Fig. 1: ntSynt pipeline overview.** The initial input for ntSynt is two or more genome assemblies. First, a Bloom filter is built which contains the *k*-mers found in all input genomes. Then, this Bloom filter is used with *indexlr*, a utility found in btllib[1], to generate ordered minimizer sketches for each input genome sequence. An undirected minimizer graph is generated from the minimizer sketches. After this graph is filtered, the initial synteny blocks are computed, followed by breaking the blocks at detected indels. The existing minimizer graph is then extended by generating additional minimizers from regions not covered by the synteny blocks using a lower minimizer window size. Next, synteny blocks are identified from this extended graph using the same filtering methods employed to generate the initial synteny blocks. This graph extension can happen for any number of decreasing window sizes. Following the graph extension rounds, collinear synteny blocks are merged to yield the final synteny blocks.

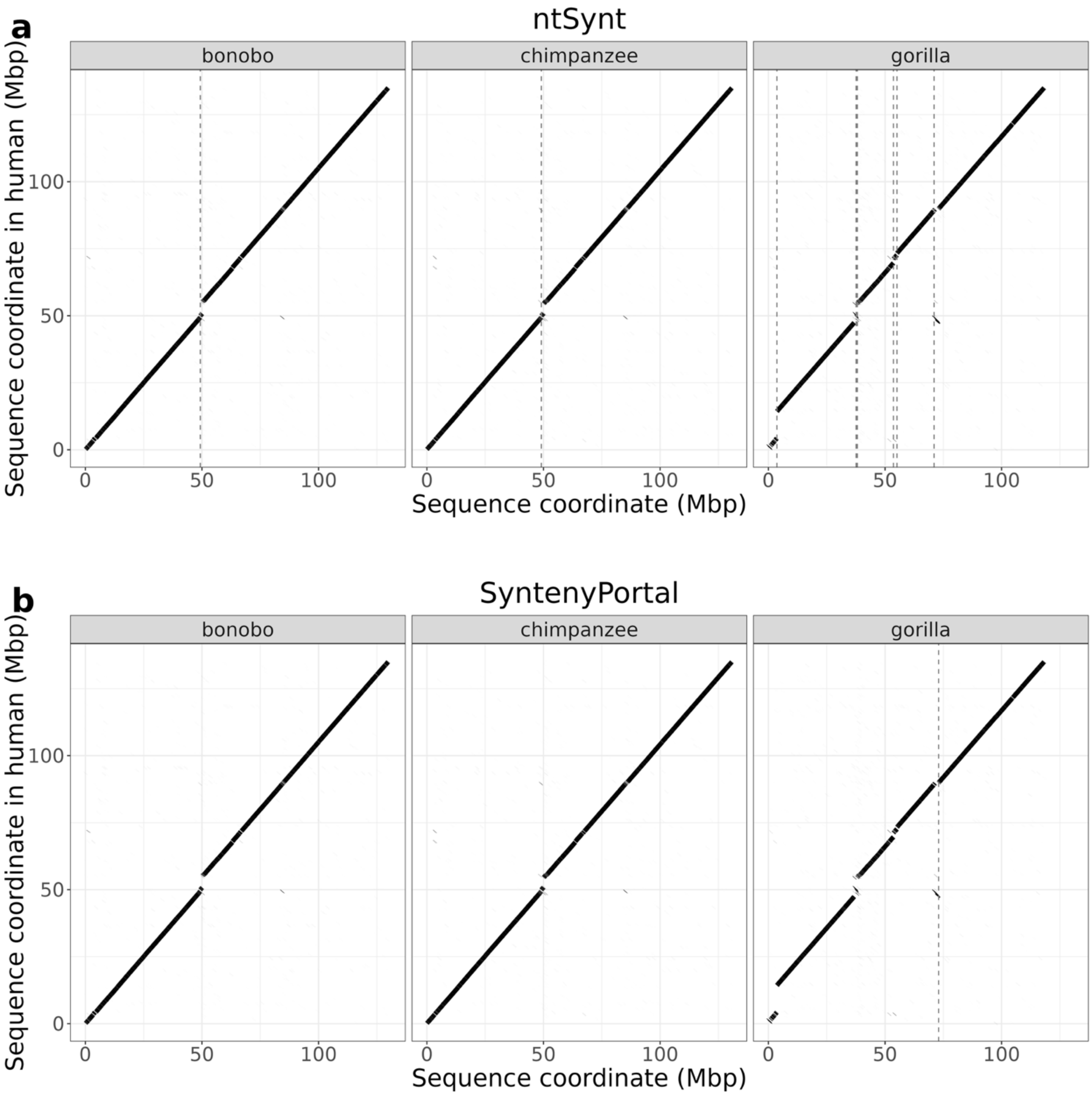

**Supplementary Fig. 2: Pairwise dot plots between human chromosome 11 and the chromosome 11 sequence of three other primate genome assemblies: bonobo, chimpanzee and gorilla.** The pairwise alignment blocks were generated using minimap2[2]. The vertical grey dashed lines denote indels greater than 300 kbp identified by (a) ntSynt or (b) SyntenyPortal[3] in any of the compared primate genomes.

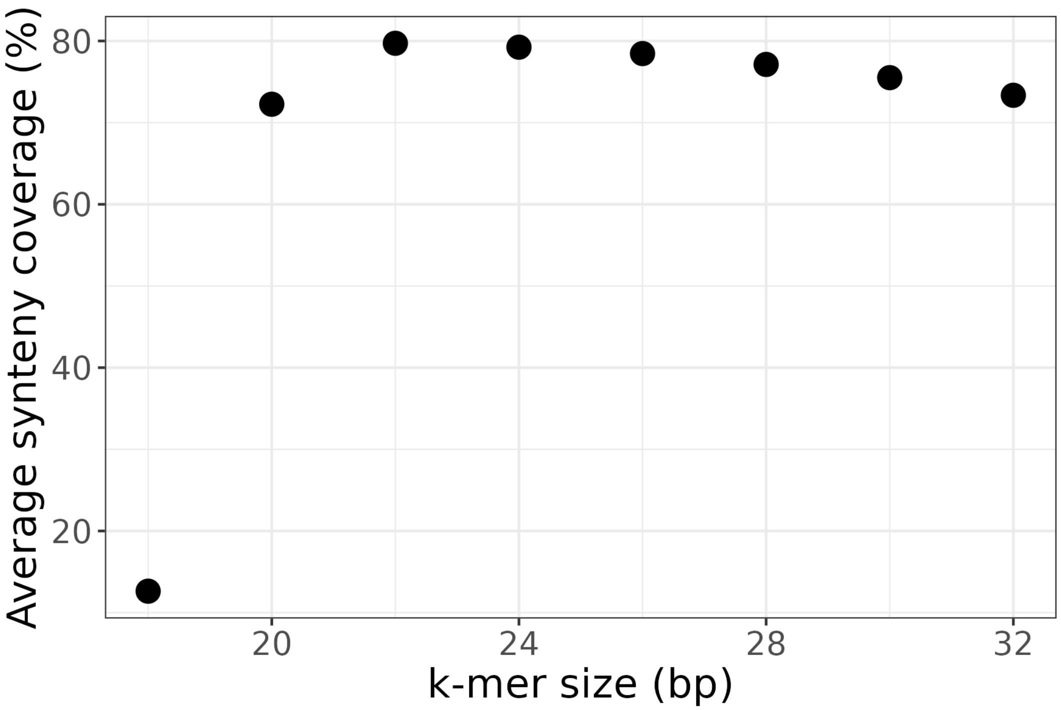

**Supplementary Fig. 3: Average synteny coverage of ntSynt blocks between human, mouse and rat reference genomes, sweeping on the *k*-mer size.** Synteny blocks were computed between the reference genomes using the parameters listed in Supplementary Table S14 with --*indel* 500000, only varying the *k*-mer size for generating the minimizer sketches.

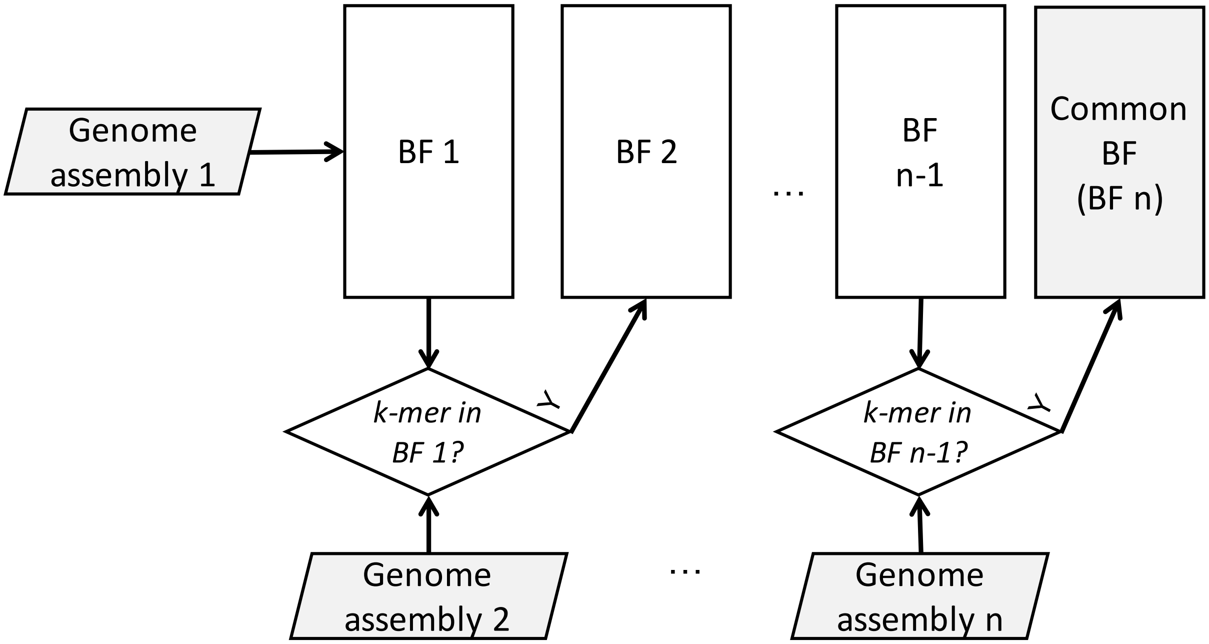

**Supplementary Fig. 4: ntSynt common Bloom filter construction using a cascading approach.** To construct the common Bloom filter (BF), which contains the *k*-mers found in each input genome assembly, the *k*-mers from the (arbitrarily chosen) initial genome assembly are first loaded into the level 1 BF (BF1). Then, the next Bloom filter (level 2, BF2) is initialized. The next input genome assembly is *k*-merized using ntHash2[4], and each *k*-mer queried against the level 1 Bloom filter. All *k*-mers that are present in the level 1 BF are added to the level 2 BF. This process continues for all input sequences, up to input genome assembly *n*, where the *k*-mers from input genome assembly *n* will be queried against BF level *n*-1, and inserted into BF level *n* if present. The level *n* BF is the final common BF, which is then used in subsequent steps. Due to the nature of the cascading approach, only two Bloom filters are kept in memory at a time.

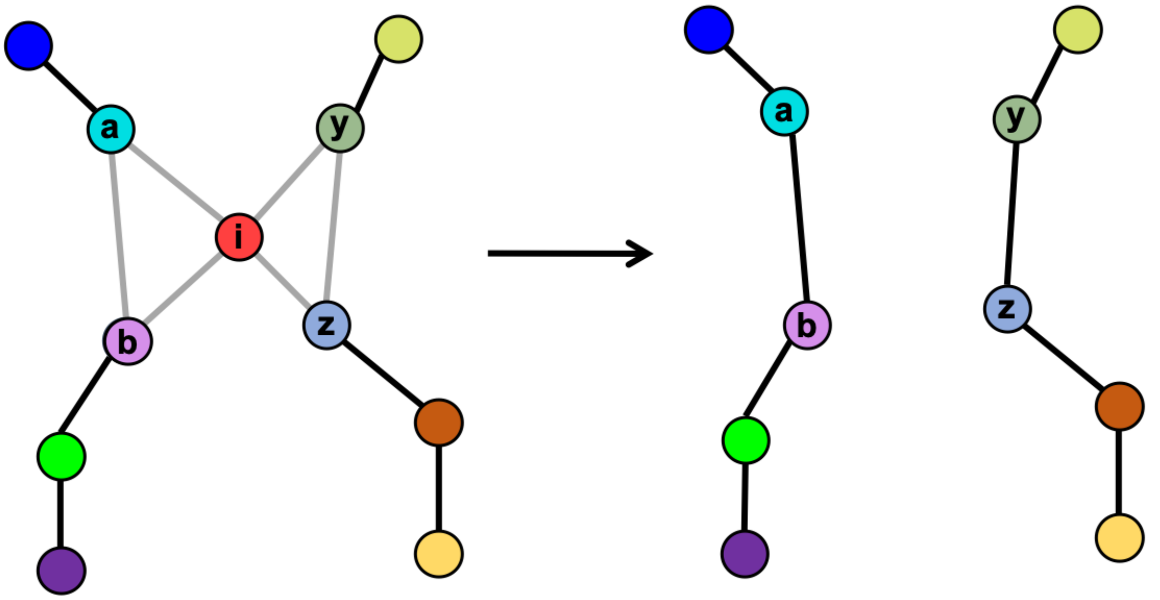

**Supplementary Fig. 5: Simplifying the ntSynt minimizer graph.** An example of the graph topology targeted by the graph simplification algorithm is shown. The nodes are minimizers, with different minimizers indicated using varying colours. The edges are coloured based on whether they have full input genome sequence input support (black; edge weight = *n*, where *n* is the number of input genomes) or if there is disagreement between genome sequence inputs (grey; edge weight < *n*). For each partially anchored edge *u, v* (degree(*u*) == degree(*v*) == 3; only one edge incident to both *u* and *v* has edge weight == *n*), the graph is traversed to find any alternate paths between *u* and *v* of length 2. For example, between the start and end nodes of partially anchored edges (*a*, *b*) and (*y*, *z*), there are alternate paths (*a, i, b*) and (*y, i, z*). For these alternate paths, the middle minimizer (*i*) is removed from the graph, and the weights of the direct edges are allocated full input sequence support.

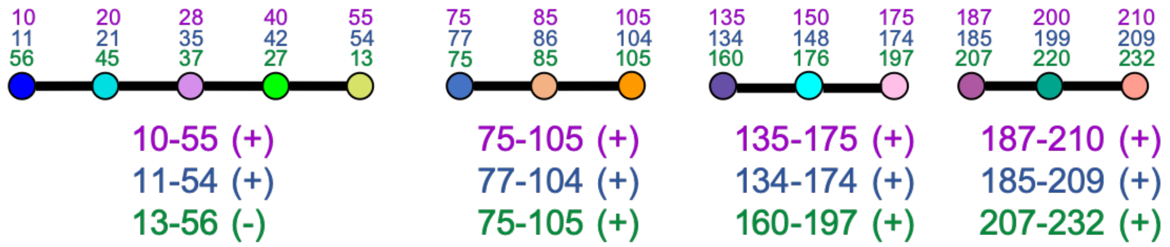

**Supplementary Fig. 6: Converting linear graph paths to synteny block coordinates.** Three input genome sequences are being compared here, indicated with purple, blue and green text. Minimizers are shown by the coloured circles, and the numbers above the minimizers indicate the position of the minimizer in the respective input sequence. The ranges and orientations below each block indicate the computed synteny block coordinates and orientations for each genome input in each synteny block.

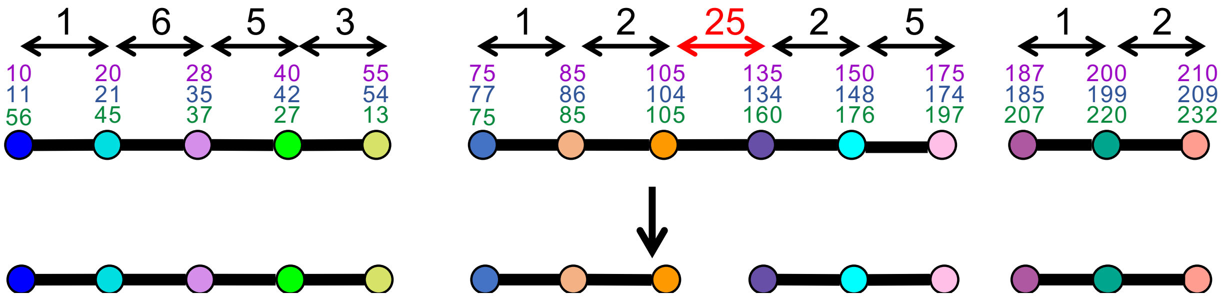

**Supplementary Fig. 7: Indel detection in ntSynt.** The positions of the given minimizer in each assembly are indicated by the coloured numbers above the minimizer graph nodes (3 different input sequences indicated by purple, blue and green). For each minimizer graph edge, the interarrival distances for each genome sequence are computed. Then, the indel score is determined from the maximum difference between these interarrival distances. If the indel score is greater than the indel score threshold (--*indel*), that edge is removed from the graph to break the synteny block at the putative indel. In the above example, the indel scores are the numbers above the double-ended arrows. For this example, assuming that the indel score threshold is set at 10, the edge indicated with the red arrow will be removed from the graph, thus breaking the synteny block at this indel.

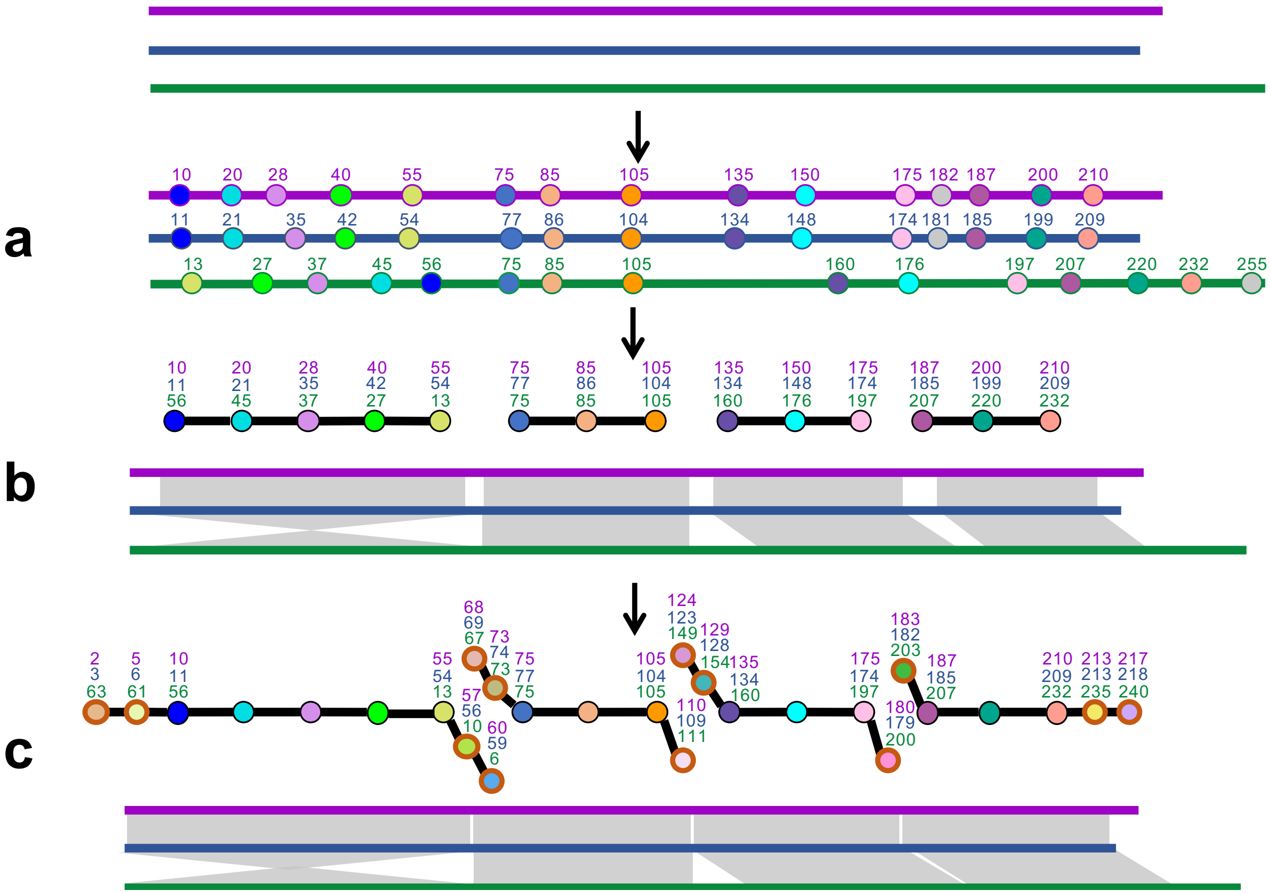

**Supplementary Fig. 8: Extension of ntSynt synteny blocks using minimizers computed with lower window sizes.** The positions of the given minimizer in each input genome sequence are indicated by the coloured numbers above the minimizer graph nodes (3 different input genome sequences indicated by purple, blue and green lines). After computing the initial synteny blocks (a-b), including indel detection, the synteny block coordinates can be refined by extending the existing minimizer graph using additional minimizers computed with a lower window size (c). Minimizers from the genomic regions that are not already covered by a synteny block are computed using a lower window size (--*w_rounds*). These minimizers are used to extend the existing graph paths, or synteny blocks, to refine their coordinates. The additional minimizers in (c) are indicated with a thick orange border.

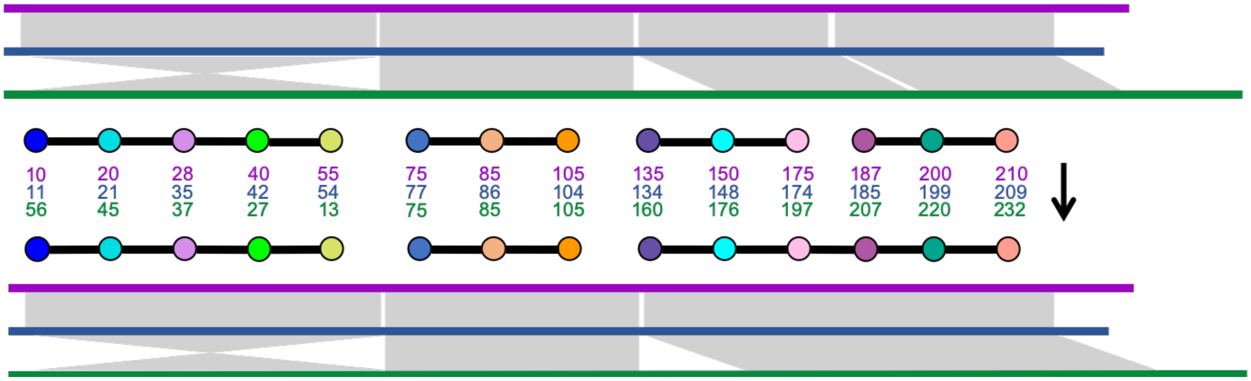

**Supplementary Fig. 9: Merging collinear synteny blocks.** Adjacent synteny blocks are considered collinear if they are not separated by an indel, are less than the merge threshold apart, and have consistent contig IDs, orientation and positions.

**Supplementary Table 1: Expected synteny block statistics for the pairwise comparisons between the human reference genome (T2T build) and one SURVIVOR[5]-simulated rearranged genome sequence, based on the ground truth.** The expected values of these statistics are the same for each rate of SNV and indels introduced with pIRS[6].

| Number of blocks | Synteny coverage (%) | Block NG50 length (Mbp) |
|---|---|---|
| 91 | 100.00 | 61.71 |

**Supplementary Table 2: Summary statistics of synteny blocks generated by ntSynt, SibeliaZ[7], halSynteny[8] and SyRI[9] using the human reference genome (T2T build) and one SURVIVOR-simulated rearranged genome.** SNVs and indels were introduced into the same SURVIVOR rearranged genome sequence at various SNV and indel rates, using pIRS. The synteny coverage and block NG50 length statistics are averaged for the two input genome sequence assemblies. The wall-clock time and memory usage statistics shown include any mapping steps required prior to running the tool (minimap2[2] for SyRI and Progressive Cactus[10] for halSynteny). The benchmarks were averaged over triplicate runs, with the average ± standard deviation tallied in the table.

| SNV rate (%) | Indel rate (%) | Tool | Number of blocks | Synteny coverage (%) | Block NG50 length (Mbp) | Wall-clock time (min) | Peak memory (GB) |
|---|---|---|---|---|---|---|---|
| 0.1 | 0.01 | **ntSynt** | 87 | 100.00 | 61.71 | 26.78 ± 4.33 | 33.99 ± 0.11 |
| | | **SibeliaZ** | 231,556 | 89.84 | 0.03 | 76.61 ± 7.71 | 31.50 ± 0.00 |
| | | **halSynteny** | 655 | 98.94 | 45.61 | 8,519.27 ± 361.45 | 17.56 ± 0.02 |
| | | **SyRI** | 93 | 100.02 | 86.30 | 19.00 ± 0.52 | 24.60 ± 0.01 |
| 1.0 | 0.10 | **ntSynt** | 129 | 99.29 | 51.73 | 26.09 ± 2.83 | 33.94 ± 0.11 |
| | | **SibeliaZ** | 142,318 | 86.93 | 0.03 | 113.34 ± 11.79 | 45.90 ± 0.02 |
| | | **halSynteny** | 879 | 99.73 | 97.22 | 8,535.16 ± 357.66 | 22.88 ± 6.90 |
| | | **SyRI** | 123 | 100.00 | 74.29 | 25.00 ± 1.05 | 23.92 ± 0.06 |
| 2.0 | 0.20 | **ntSynt** | 226 | 96.90 | 44.30 | 37.37 ± 2.39 | 33.90 ± 0.11 |
| | | **SibeliaZ** | 172,172 | 85.74 | 0.03 | 147.06 ± 14.82 | 54.69 ± 0.00 |
| | | **halSynteny** | 718 | 99.77 | 92.60 | 8,517.66 ± 345.14 | 29.74 ± 2.11 |
| | | **SyRI** | 151 | 99.97 | 84.34 | 32.14 ± 1.07 | 22.30 ± 0.03 |
| 3.0 | 0.30 | **ntSynt** | 269 | 95.44 | 39.89 | 49.94 ± 8.77 | 33.91 ± 0.12 |
| | | **SibeliaZ** | 234,919 | 83.88 | 0.02 | 195.36 ± 20.32 | 58.68 ± 0.00 |
| | | **halSynteny** | 676 | 99.58 | 94.57 | 8,524.84 ± 346.54 | 35.04 ± 0.13 |
| | | **SyRI** | 218 | 99.92 | 74.29 | 37.80 ± 1.52 | 21.70 ± 0.02 |
| 4.0 | 0.40 | **ntSynt** | 277 | 94.80 | 43.21 | 61.49 ± 13.14 | 33.83 ± 0.23 |
| | | **SibeliaZ** | 358,553 | 80.73 | 0.01 | 267.79 ± 12.64 | 59.62 ± 0.00 |
| | | **halSynteny** | 461 | 98.80 | 96.69 | 8,511.31 ± 357.38 | 44.02 ± 0.80 |
| | | **SyRI** | 164 | 99.97 | 74.29 | 61.31 ± 2.55 | 29.35 ± 0.05 |
| 5.0 | 0.50 | **ntSynt** | 302 | 94.25 | 35.91 | 72.17 ± 13.82 | 33.91 ± 0.17 |
| | | **SibeliaZ** | 545,068 | 74.39 | 0.00 | 339.58 ± 33.23 | 58.68 ± 0.01 |
| | | **halSynteny** | 510 | 99.63 | 97.68 | 8,528.08 ± 359.78 | 55.81 ± 4.53 |
| | | **SyRI** | 208 | 99.95 | 74.29 | 64.64 ± 2.24 | 27.54 ± 0.09 |

**Supplementary Table 3: Analysis of the genomic regions that are not covered by ntSynt synteny blocks when comparing the human reference genome (T2T build) to one SURVIVOR-rearranged genome sequence assembly with different variant (SNV+indel) rates.** The regions of the reference genome sequence that were not covered by synteny blocks were compared with annotated centromere coordinates.

| SNV rate (%) | Indel rate (%) | Centromeric? | Percentage of genome without synteny block coverage (%) |
|---|---|---|---|
| 0.1 | 0.01 | No | 0.03 |
| | | Yes | 0.00 |
| 1.0 | 0.10 | No | 0.03 |
| | | Yes | 0.71 |
| 2.0 | 0.20 | No | 0.04 |
| | | Yes | 3.08 |
| 3.0 | 0.30 | No | 0.16 |
| | | Yes | 4.43 |
| 4.0 | 0.40 | No | 0.19 |
| | | Yes | 5.03 |
| 5.0 | 0.50 | No | 0.27 |
| | | Yes | 5.50 |

**Supplementary Table 4: Expected synteny block statistics for the multi-genome comparisons between the human reference genome (T2T build) and four SURVIVOR-simulated rearranged genomes, based on the ground truth.** The expected values of the statistics are the same for each rate of SNV and indels introduced using pIRS.

| Number of blocks | Synteny coverage (%) | Block NG50 length (Mbp) |
|---|---|---|
| 295 | 100.00 | 24.59 |

**Supplementary Table 5: Summary statistics of synteny blocks generated by ntSynt and SibeliaZ using the human reference genome (T2T build) and four SURVIVOR-simulated rearranged genomes.** SNVs and indels were introduced into the SURVIVOR rearranged genomes at various rates using pIRS. The synteny coverage and block NG50 length statistics are averaged for the five input genome sequence assemblies. For the block NG50 length statistics, "-" denotes where this statistic could not be calculated due the synteny coverage being less than half of the input genome size. The benchmarks were averaged over triplicate runs, with the average ± standard deviation tallied in the table.

| Tool | SNV rate (%) | Indel rate (%) | Number of blocks | Synteny coverage (%) | Block NG50 length (Mbp) | Wall-clock time (h) | Peak memory (GB) |
|---|---|---|---|---|---|---|---|
| **ntSynt** | 0.1 | 0.01 | 290 | 99.87 | 24.59 | 1.03 ± 0.29 | 34.05 ± 0.02 |
| **ntSynt** | 1.0 | 0.10 | 315 | 98.67 | 23.73 | 1.25 ± 0.54 | 34.01 ± 0.06 |
| **ntSynt** | 2.0 | 0.20 | 384 | 95.26 | 21.97 | 1.23 ± 0.53 | 33.95 ± 0.05 |
| **ntSynt** | 3.0 | 0.30 | 466 | 93.24 | 20.34 | 1.33 ± 0.60 | 34.01 ± 0.05 |
| **ntSynt** | 4.0 | 0.40 | 485 | 92.12 | 18.36 | 1.29 ± 0.63 | 33.98 ± 0.02 |
| **ntSynt** | 5.0 | 0.50 | 477 | 91.11 | 16.78 | 1.28 ± 0.54 | 34.02 ± 0.04 |
| **SibeliaZ** | 0.1 | 0.01 | 163,945 | 90.78 | 0.05 | 2.01 ± 0.29 | 64.46 ± 0.02 |
| **SibeliaZ** | 1.0 | 0.10 | 169,377 | 85.98 | 0.03 | 4.67 ± 1.06 | 142.06 ± 0.06 |
| **SibeliaZ** | 2.0 | 0.20 | 392,391 | 76.28 | 0.01 | 9.44 ± 1.80 | 179.87 ± 0.04 |
| **SibeliaZ** | 3.0 | 0.30 | 557,286 | 45.94 | - | 20.45 ± 4.19 | 194.17 ± 0.01 |
| **SibeliaZ** | 4.0 | 0.40 | 159,549 | 9.50 | - | 39.06 ± 7.18 | 195.58 ± 0.00 |
| **SibeliaZ** | 5.0 | 0.50 | 20,947 | 0.76 | - | 58.17 ± 13.11 | 189.70 ± 0.02 |

**Supplementary Table 6: Reference genome assemblies used for synteny block analysis with ntSynt, SibeliaZ and SyntenyPortal.**

| Species | Common name | Build | Accession |
|---|---|---|---|
| *Homo sapiens* | Human | GRCh38 | GCA_000001405.15 |
| *Pan paniscus* | Bonobo | Mhudiblu_PPA_v0 | GCF_013052645.1 |
| *Pan troglodytes* | Chimpanzee | Clint_PTRv2 | GCF_002880755.1 |
| *Gorilla gorilla gorilla* | Gorilla | Kamilah_GGO_v0 | GCF_008122165.1 |
| *Mus musculus* | Mouse | GRCm39 | GCA_000001635.9 |
| *Rattus norvegicus* | Rat | Rnor_6.0 | GCA_000001895.4 |

**Supplementary Table 7: Contiguity, coverage and benchmarking statistics for synteny blocks computed on human, bonobo, chimpanzee and gorilla reference genome builds using ntSynt, SibeliaZ, and the SyntenyPortal web application.** Synteny coverage of complete blocks refers to synteny blocks that include all four input genome assemblies. SyntenyPortal is a web application based on pre-computed sequence alignments, therefore benchmarks cannot be computed for this resource.

| Tool | Number of synteny blocks | Synteny coverage (%) | Synteny coverage of complete blocks (%) | NG50 block length (Mbp) | Wall-clock time (h) | Peak memory (GB) |
|---|---|---|---|---|---|---|
| **ntSynt** | 1,951 | 92.23 | 92.23 | 7.61 | 0.80 | 32.19 |
| **SibeliaZ** | 136,961 | 88.63 | 85.35 | 0.05 | 2.71 | 87.67 |
| **SyntenyPortal** | 186 | 94.69 | 94.69 | 48.14 | N/A | N/A |

**Supplementary Table 8: Mapped read statistics from aligning gorilla PacBio HiFi reads (individual Kamilah) to the gorilla reference genome build used in the synteny tests (Kamilah_GGO_v0) and a newer gorilla reference genome build (mGorGor1).** All HiFi sequencing reads were aligned to the human genome reference sequence (GRCh38) using minimap2, and the reads that aligned to a putative deletion in the gorilla genome (Kamilah_GGO_v0 chromosome 11, coordinates 4,249,395-14,277,464) were extracted using samtools[11]. The extracted reads were then aligned to the two gorilla genome builds using minimap2, and filtered to retain primary mapped reads with mapping quality >= 50. The PacBio HiFi reads are available from SRA under accession SRR13446351 and the mGorGor1 assembly is available from GenBank under accession GCA_029281585.1.

| Assembly build | Total number of reads | Number of mapped reads |
|---|---|---|
| **Kamilah_GGO_v0** | 7,212 | 185 (2.6%) |
| **mGorGor1** | 7,212 | 7,207 (99.9%) |

**Supplementary Table 9: Genome sequence assemblies used for synteny block analysis with ntSynt and SibeliaZ.**

| Species | Common name | Cell line /individual | Assembler | Accession |
|---|---|---|---|---|
| *Homo sapiens* | Human | NA24385 | GoldRush[12] | https://doi.org/10.5281/zenodo.7884681 |
| *Pan paniscus* | Bonobo | Mhudiblu | hifiasm[13] | GCA_030221875.1 |
| *Pan troglodytes* | Chimpanzee | Clint | hifiasm | GCA_030128855.1 |
| *Gorilla gorilla gorilla* | Gorilla | Kamilah | hifiasm | GCA_030174185.1 |

**Supplementary Table 10: Contiguity, coverage and benchmarking statistics for synteny blocks computed between human, bonobo, chimpanzee and gorilla genome assemblies using ntSynt and SibeliaZ.** Synteny coverage of complete blocks refers to synteny blocks that include all four input genome sequence assemblies. The human genome assembly was generated using GoldRush, while the other assemblies were produced using hifiasm.

| Tool | Number of synteny blocks | Synteny coverage (%) | Synteny coverage of complete blocks (%) | NG50 block length (Mbp) | Wall-clock time (h) | Peak memory (GB) |
|---|---|---|---|---|---|---|
| **ntSynt** | 7,450 | 82.81 | 82.81 | 1.30 | 1.18 | 32.76 |
| **SibeliaZ** | 145,678 | 83.29 | 79.13 | 0.05 | 3.24 | 102.64 |

**Supplementary Table 11: Contiguity, coverage and benchmarking statistics for synteny blocks between human, mouse and rat reference genome assemblies using ntSynt and the SyntenyPortal web application.** Synteny coverage of complete blocks refers to synteny blocks that include all three input genome assemblies. SyntenyPortal is a web application based on pre-computed sequence alignments, therefore benchmarks cannot be computed for this resource.

| Tool | Number of synteny blocks | Synteny coverage (%) | Synteny coverage of complete blocks (%) | NG50 block length (Mbp) | Wall-clock time (h) | Peak memory (GB) |
|---|---|---|---|---|---|---|
| **ntSynt** | 1,069 | 79.24 | 79.24 | 3.86 | 1.08 | 34.22 |
| **SyntenyPortal** | 806 | 89.79 | 89.79 | 7.05 | N/A | N/A |

**Supplementary Table 12: Breakdown of benchmarking statistics for computing synteny blocks between human, mouse and rat reference genomes using ntSynt.** The ntSynt stages were grouped into three main steps. Note that some processes within the minimizer computation step can run concurrently, thus the sum of times over all steps will be greater than the total wall-clock time of the entire process.

| ntSynt step | Wall-clock time (min) | Peak memory (GB) |
|---|---|---|
| **common BF construction** | 5.38 | 34.22 |
| **minimizer computation** | 18.98 | 18.13 |
| **synteny block construction** | 46.95 | 25.31 |

**Supplementary Table 13: Output format of ntSynt synteny blocks file.** The output file is in tab-separated format.

| Column number | Description |
|---|---|
| **1** | Synteny block ID - Lines with the same ID are part of the same synteny block |
| **2** | Genome file name |
| **3** | Genome chromosome/contig |
| **4** | Genome start coordinate |
| **5** | Genome end coordinate |
| **6** | Strand |
| **7** | Number of mapped minimizers in this synteny block |
| **8** | Reason for discontinuity with previous synteny block |

**Supplementary Table 14: Default ntSynt parameter settings based on the user-supplied divergence (--*divergence*).**

| Divergence range | Presets (default parameters) |
|---|---|
| < 1% | --block_size 500 --indel 10000 --merge 10000 --w_rounds 100 10 |
| 1% - 10% | --block_size 1000 --indel 50000 --merge 100000 --w_rounds 250 100 |
| >10% | --block_size 10000 --indel 100000 --merge 1000000 --w_rounds 500 250 |

**Supplementary Table 15: Parameter settings used when running synteny block comparator tools.**

| Tool | Sequence divergence | Parameters |
|---|---|---|
| SibeliaZ (+ maf2synteny) | < 1% | *SibeliaZ*: -n -t12<br>*maf2synteny*: -b 500 |
| | >= 1% | *SibeliaZ*: -n -t12<br>*maf2synteny*: -b 1000 |
| halSynteny (+ Progressive Cactus) | < 1% | *Progressive Cactus*: --maxCores 12 --binariesMode local<br>*halSynteny*: --minBlockSize 500 --maxAnchorDistance 10000 |
| | >= 1% | *Progressive Cactus*: --maxCores 12 --binariesMode local<br>*halSynteny*: --minBlockSize 1000 --maxAnchorDistance 100000 |
| SyRI (+ minimap2) | < 1% | *minimap2*: -x asm -t 12 -c --eqx<br>*SyRI*: --nosnp --nc 12 -F P --invgaplen 10000 --tdgaplen 10000<br>--no-chrmatch |
| | >= 1% | *minimap2*: -x asm -t 12 -c --eqx<br>*SyRI*: --nosnp --nc 12 -F P --invgaplen 100000 --tdgaplen 100000<br>--no-chrmatch |

**Supplementary References**

1. Nikolić, V. *et al.* btllib: A C++ library with Python interface for efficient genomic sequence processing. *Journal of Open Source Software* **7**, 4720 (2022).
2. Li, H. Minimap2: pairwise alignment for nucleotide sequences. *Bioinformatics* **34**, 3094–3100 (2018).
3. Lee, J. *et al.* Synteny Portal: a web-based application portal for synteny block analysis. *Nucleic Acids Research* **44**, W35–W40 (2016).
4. Kazemi, P. *et al.* ntHash2: recursive spaced seed hashing for nucleotide sequences. *Bioinformatics* **38**, 4812–4813 (2022).
5. Jeffares, D. C. *et al.* Transient structural variations have strong effects on quantitative traits and reproductive isolation in fission yeast. *Nature Communications* **8**, 14061 (2017).
6. Hu, X. *et al.* pIRS: Profile-based Illumina pair-end reads simulator. *Bioinformatics* **28**, 1533–1535 (2012).
7. Minkin, I. & Medvedev, P. Scalable multiple whole-genome alignment and locally collinear block construction with SibeliaZ. *Nature Communications* **11**, 6327 (2020).
8. Krasheninnikova, K. *et al.* halSynteny: a fast, easy-to-use conserved synteny block construction method for multiple whole-genome alignments. *GigaScience* **9**, giaa047 (2020).
9. Goel, M., Sun, H., Jiao, W.-B. & Schneeberger, K. SyRI: finding genomic rearrangements and local sequence differences from whole-genome assemblies. *Genome Biology* **20**, 277 (2019).
10. Armstrong, J. *et al.* Progressive Cactus is a multiple-genome aligner for the thousand-genome era. *Nature* **587**, 246–251 (2020).
11. Danecek, P. *et al.* Twelve years of SAMtools and BCFtools. *Gigascience* **10**, (2021).
12. Wong, J. *et al.* Linear time complexity de novo long read genome assembly with GoldRush. *Nature Communications* **14**, 2906 (2023).
13. Cheng, H., Concepcion, G. T., Feng, X., Zhang, H. & Li, H. Haplotype-resolved de novo assembly using phased assembly graphs with hifiasm. *Nature Methods* **18**, 170–175 (2021).
